# scROSHI - robust supervised hierarchical identification of single cells

**DOI:** 10.1101/2022.04.05.487176

**Authors:** Michael Prummer, Anne Bertolini, Lars Bosshard, Florian Barkmann, Josephine Yates, Valentina Boeva, The TumorProfiler Consortium, Daniel Stekhoven, Franziska Singer

**Affiliations:** Nexus Personalized Health Technologies, ETH Zurich, Zurich Switzerland; Institute for Machine Learning, Department of Computer Science, ETH Zurich, Zurich Switzerland; Cochin Institute, Inserm U1016, CNRS UMR 8104, Paris Descartes University UMR-S1016, Paris 75014, France; Swiss Institute of Bioinformatics (SIB), Zurich, Switzerland

## Abstract

Identifying cell types based on expression profiles is a pillar of single cell analysis. Existing machine-learning methods identify predictive features from annotated training data, which are often not available in early-stage studies. This can lead to overfitting and inferior performance when applied to new data. To address these challenges we present scROSHI, which utilizes previously obtained cell type-specific gene sets and does not require training or the existence of annotated data. By respecting the hierarchical nature of cell type relationships and assigning cells consecutively to more specialized identities, excellent prediction performance is achieved. In a benchmark based on publicly available PBMC data sets, scROSHI outperforms competing methods when training data are limited or the diversity between experiments is large.

## Introduction

After more than two decades of technological development from its earliest attempts^1,2^, single cell transcriptomics studies have come of age and are widely used for basic as well as translational research^3–5^. This is best showcased by the recent explosion of single cell atlases of various organs and organisms^6–9^, as well as the use of single cell transcriptomics for disease investigation^10^. The term “atlas’’ describes the result of identifying each and every cell type in the analyzed tissue sample for known cell types and discovering novel cell types defined by their transcriptomic phenotype. In the early days, this was often a labor-intensive manual process requiring expert field knowledge, and for unknown or novel cell types such as developing or precursor cells it still is.

In recent years, a large number of tools have been developed to automate cell type identification with varying performance, as summarized in a recent benchmark study^11^. The common theme of these tools is that the expression profile of a target cell is compared to known expression profiles of particular cell types, possibly limited to a subset of genes that are relatively stable and highly expressed, so-called marker genes. In order to derive expression features predictive for a cell type, it is commonplace to use unsupervised clustering of a single cell data set and assign the cluster labels to cell types based on biological interpretation. In the next step, this cell type label is interpreted as the ground truth to build a machine learning model which finds the features relevant for cell type prediction.

However, as will be shown in the course of this work, learning features and performing the classification on the same data can lead to overfitting even if separate training and test data are used, provided they both were acquired under the same experimental condition. As a consequence, the cell type classification uncertainty is underestimated. Another challenge in cell type classification is that sometimes the number of possible candidate cell types is large, which tends to increase the misclassification rate.

To address these challenges, we present **s**ingle **c**ell **Ro**bust **S**upervised **H**ierarchical **I**dentification of cell types (scROSHI), which utilizes a-priori defined cell type-specific gene sets and does not require training or the existence of annotated data. By respecting the hierarchical nature of cell type relationships due to differentiation within a lineage and assigning cells consecutively to more specialized identities, excellent prediction performance is achieved. We provide evidence for this by comparing the performance of scROSHI with three existing tools that scored among the best in a recent benchmark study^11^. There, the authors used the same datasets for training, testing, and validation. Here, based on three annotated datasets, we show that scROSHI outperforms the competing methods when applied to data from a different source than the training data, which is a much more realistic scenario.

Taken together, scROSHI is a conceptually simple, transparent, interpretable, and robust cell type classification approach particularly useful when previous knowledge about cell type-specific genes is available but annotated training data is scarce. scROSHI is available as an R package and can thus be seamlessly integrated into single cell analysis workflows.

## Methods

### scROSHI: design considerations

The most important design criterion for scROSHI was that the method should be capable of automated classification in the absence of labeled training data. This excluded any machine learning approach that would require training a model. Instead, the method should utilize and rely on the vast amount of validated cell type-specific gene sets available from previous bulk or single cell experiments, such as the widely acknowledged immune cell type gene sets (also known as “*lm22*”) used by the cibersort algorithm^12^ or the somewhat related resource for single cell melanoma data^13^.

Another important design criterion was that the method should avoid re-training on new datasets to avoid overfitting. Provided the originally chosen gene set was previously validated to be robust against changes in the experimental condition, re-training is not necessary.

This argument can be turned around to provide a strategy on how to arrive at a suitable gene set for cell types for which a previously validated set is not available: included as marker genes, i.e., genes that are highly expressed in the target cell type, should be those genes that have this property independent of tissue type, sample type (i.e., cultured immortalized cells, cultured primary cells, tissue biopsy), detailed setup (culture medium, organism), or patient characteristics (gender, ethnicity, age). The larger the diversity of the test data, the more robust and broadly applicable the final gene set.

Given a set of cell type-specific genes and without the need for training a model, one possibility to assign a cell type from a list of candidates to a target cell is to test for association and choose the one that fits best. On one end of the spectrum when measuring association is the hypergeometric test comparing the proportion of highly expressed genes specific for one cell type with the proportion of highly expressed genes specific for all other cell types. The advantage is that it can almost always be calculated and is robust against expression outliers. On the one hand, it is simple to use because the cell type-specific reference does not have to be known quantitatively in the form of an expression profile. On the other hand, it is relatively insensitive because it completely ignores the quantitative nature of the expression profile of the target cell, which is typically available. On the other end of the spectrum when measuring association one can quantitatively match the expression profile of a known cell type to the expression profile of the target cell, for instance, using Spearman’s correlation. While this approach is robust against expression outliers, it is relatively expensive in data availability, i.e., it requires knowledge of the gene expression profile of the reference. We chose to follow an intermediate path by performing a quantitative and robust test, the Mann-Whitney rank sum test, to compare the expression ranks of the genes specific for one cell type with the ranks of the genes specific for all other cell types. The negative log of the test’s p-value is then a measure of the association strength between the target cell and this cell type.

Similarly, as in most classification problems, it is assumed here that a cell belongs to one and only one of a given set of classes. However, scROSHI goes one step further and allows for the introduction of two additional classes, *unknown* and *uncertain*, to deal with the unavoidable classification uncertainty.

Another step to improve on the classification efficiency is to utilize the hierarchical tree structure that is inherent to cell types due to developmental specialization and therefore apply a hierarchical classification approach. Instead of classifying all cell types at once, target cells are first assigned to a smaller number of major cell types and then consecutively to more specialized classes. This way, relatively similar cell types can be distinguished provided they belong to different branches in the tree.

### The scROSHI workflow

1. Find out which cell types to expect from field knowledge.
2. Obtain validated cell type-specific gene sets from the literature or learn marker genes based on other datasets.
3. Obtain a hierarchical tree structure to define cell type parent - kin relationships (“generations”).
4. For each cell and each cell type in the first generation, determine the p-value of a one-sided Mann-Whitney test of the Null hypothesis that the expression rank sum of the genes specific for this cell type is the same as or smaller than the rank sum of the genes specific for any other cell type in the list. The alternative hypothesis is that the rank sum of the genes specific for this cell type is larger.
5. Compute the normalized negative log of the p-value for each cell and each cell type, respectively. Interpret the result as the probability of the cell belonging to that cell type.
6. If none of the probabilities is above a certain threshold, do not assign a cell type label to the cell but assign it to the class “unknown”.
7. If the ratio between the largest and the second largest probability is below a certain threshold, do not assign a cell type label to the cell but assign it to the class “uncertain”.
8. Repeat 4 to 7 for the second generation, and so on. For a certain parent cell type to be further classified into next generation cell types, also include uncertain or unknown cells with high similarity to this parent cell type as defined by knn classification.

scROSHI takes as input the gene x cell count matrix, either raw or normalized (Fig. 1A). The choice of normalization and/or transformation method has little influence on the results because the cell type score is based on ranks rather than on the actual values. In our studies we typically use sctransform^14^, which corrects unwanted biases using regularized negative binomial regression.

**Figure 1:**
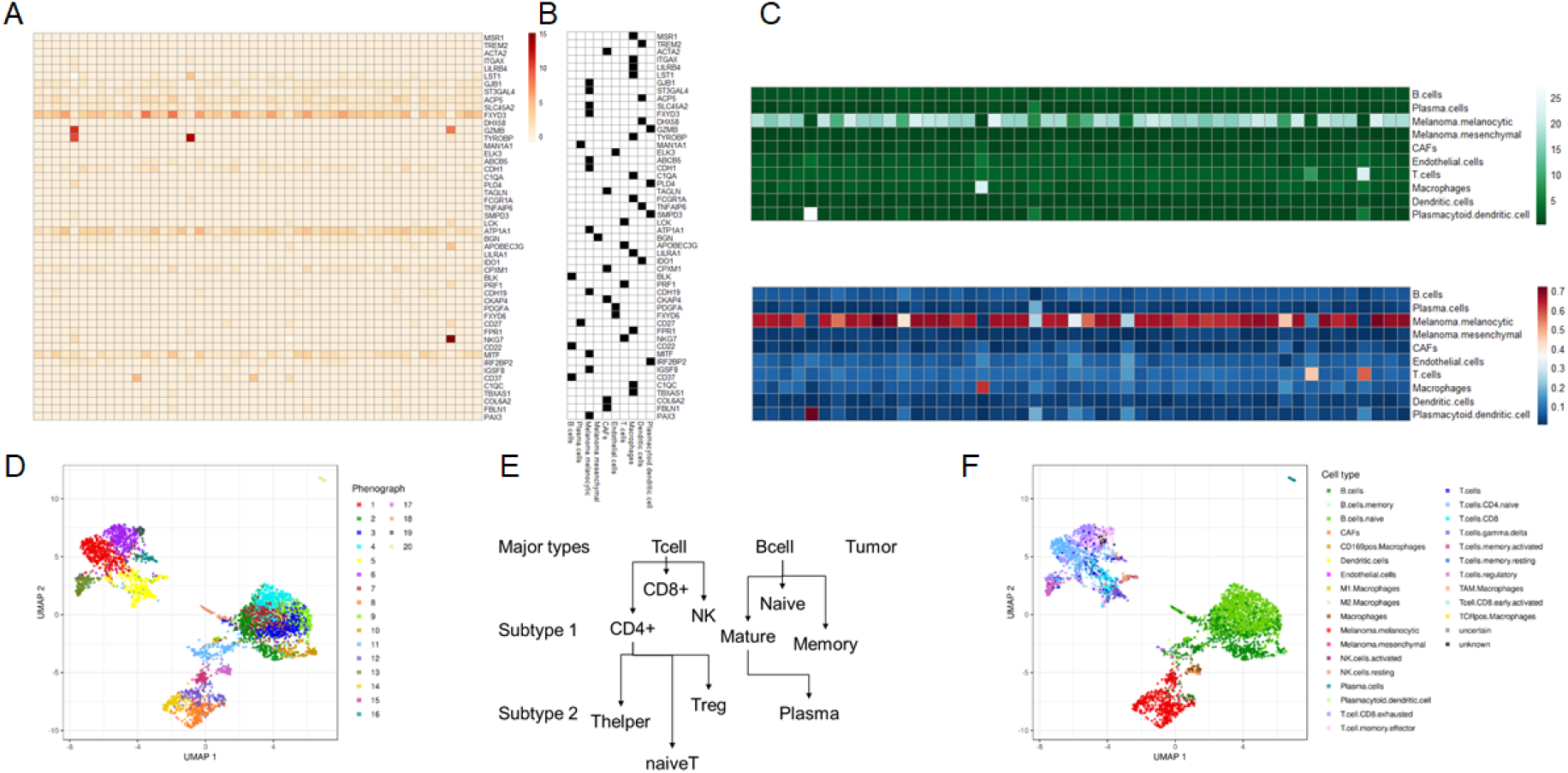
Schematics of the scROSHI workflow. The gene x cell normalized expression matrix (A) is combined with the binary gene x cell type membership matrix (B) to define genes specific for a cell type (black) and genes specific for other cell types (white). (C) The pair-wise one-sided Mann-Whitney rank sum test provides a cell type x cell score matrix (top), which is normalized to give probabilities (bottom). (D) UMAP representation of all cells from a melanoma patient biopsy using the most highly variable genes, colored by phenograph clusters. (E) Developmental “family tree” defining cell type hierarchies. (F) The representation in D is colored by scROSHI predicted cell types.

The second ingredient required for cell type classification is a collection of cell type-specific gene sets (Fig. 1B). The selection of cell types to expect will depend on the nature of the sample. It is recommended to adapt the cell type selection to keep classification specificity high whenever possible: having closely related cell types in the candidate list may sometimes be required but should be avoided if possible.

There are a number of resources available containing curated reference datasets, mostly assembled from bulk RNA-seq or microarray data of sorted cell types. Examples are the C8 set of the MSigDB collection^15^, the lm22 immune cell list of cibersort^12^, the BioGPS Human Cell Type and Tissue Gene Expression Profiles collection^16^ from harmonizome^17^, or the Bioconductor^18^ package celldex^19^. These references are often good enough for most applications provided that they contain the cell types that are expected to be present in the data at hand. For our contribution to the Tumor Profiler Study^20^, working on melanoma patient biopsy samples, we used the curated gene set from Tirosh et al.^13^ in combination with the immune cell list of cibersort.

In cases where quantitative cell type-specific reference profiles are available they can be used as is or they can be binarized to obtain cell type-specific gene lists (Fig. 1B). They should contain genes that show little variability and are highly expressed in the target cell type and have zero or weak expression in all other cell types. The gene lists do not need to be exclusive, i.e., the same gene can appear in different cell type lists, but the overlap between cell types should be kept small. Obviously, the more similar two cell types are, the larger the overlap between their specific gene lists will be. In addition, given the sparsity of single cell count data, gene lists with only a few members will not have enough sensitivity compared to larger lists.

The third ingredient to scROSHI is a hierarchical tree structure defining parent - kin relationships between cell types (generations). The purpose of this tree is to classify cells first into a small number of coarse-grained cell type super-families and then consecutively into more and more specialized, fine-grained cell (sub-)types (Fig. 1E). This way, the number of possible candidate cell types in each step is much smaller than the total number of candidate cell types thus reducing the possibility of false classification.

Based on these inputs, scROSHI performs the cell type score assignment and classification. For each cell and each cell type, a one-sided Mann-Whitney U-test is performed. The Null hypothesis is that the expression rank sum of the genes specific for this cell type is the same as or smaller than the rank sum of the genes specific for any other cell type in the list. The alternative hypothesis is that the rank sum of the genes specific for this cell type is larger. The normalized negative log of the p-value for each cell-cell type pair is interpreted as the probability of the cell belonging to the respective cell type (Fig. 1C). If none of the probabilities is above a certain threshold, no cell type label is assigned to the cell but the class “unknown”. Also, if the ratio between the largest and the second largest probability is below a certain threshold, again no cell type label is assigned to the cell but the class “uncertain”. Both categories, “unknown” and “uncertain”, can reflect populations that lack the proper cell type in the list of *a priori* selected cell types.

Taken together, these steps facilitate an enrichment of the pure data-driven description of the single-cell data (Fig. 1D) with biological meaning (Fig. 1F).

### Benchmarking

A detailed description of the datasets used, their origin, which preprocessing steps were applied, as well as the description of the pipeline and the competing tools is described in the Supplementary Material.

To briefly summarize, we used public datasets with a similar cell type composition to benchmark scROSHI against high profile competitor methods. Data from three peripheral blood mononuclear cell experiments were retrieved, one from an adult human in which the cell types were pre-sorted (Zheng_sorted set), and one each of an adult (Adult set) and a newborn (Newborn set). Hence, the three sets are similar in content but differ in experimental setting and donor age.

We defined a common set of matching cell type labels across the three datasets for comparisons between datasets (see Supplementary Material for further details on the ground truth dictionary).

Based on a previous benchmark of automatic cell identification methods ^11^, we decided to compare scROSHI to three front runners: support vector machine (SVM), random forest (RF) and GARNETT^21^. The main difference between scROSHI and its competitors is the fact that they use part of the data to train a model whereas with scROSHI there is no training involved once the cell type-specific gene sets are selected. While SVM and RF can capture non-linear relationships between the explanatory features (gene expression) as well as interactions between them, GARNETT is based on a penalized multivariate generalized linear prediction model (GLMNET).

The competing methods were used under standard conditions, with default parameter settings.

We trained a model for RF, SVM, and GARNETT and evaluated the performance of the classifiers by applying a 5-fold cross-validation for each dataset. The folds were split in a stratified manner in order to keep equal proportions of each cell population in each fold. We used the same training and testing folds for all classifiers. scROSHI and Garnett required a marker gene list. We used a list of marker genes based on previous publications^12,22^ for scROSHI and a garnett-optimized marker list (`check_markers()` function from the garnett package v.0.2.17) for Garnett..

### Validation scheme

Each of the three datasets was split in training, validation, and testing sets. Three major validation runs were performed in which each of the three datasets served as the training/validation set. After the final model was obtained, it was tested once “in set” on the testing set that came from the same experiment as the training data, and two times “out of set” on the two remaining sets from which the model has not yet seen any data. Further details on the validation scheme can be found in the Supplementary Material.

### Copy number variation estimation

To pre-process scRNA-seq data from the Tumor Profiler Study, we used a procedure based on standard quality control measures^23^. First, to retain only high quality cells, we removed cells with fewer than 700 expressed genes and 1,500 total read counts detected. Second, to avoid contamination by dying cells while retaining as many informative cells as possible, we filtered out cells with more than 35% of read counts coming from mitochondrial genes^24,25^.

To distinguish normal from malignant cells, we inferred large-scale copy number variations (CNVs) from the gene expression data using *infercnvpy*^*26*^. We ran *infercnvpy* on every sample individually using T cells, B cells, Endothelial cells and Macrophages as reference cells. The gene ordering file containing the chromosomal start and end position for each gene was generated from the human GRCh37 assembly. To reduce the noise level, we only used genes that had a mean read count greater than 0.1.

We then used an approach based on hierarchical clustering of single cell copy number profiles to detect cells with and without CNVs. After calling CNVs, we used *scipy*’s implementation of hierarchical clustering with Ward linkage^27^ to obtain a dendrogram of the CNV profiles. By definition, each node in a dendrogram only had two child nodes that represented a cluster of clusters, except for leaf nodes that represented a cluster of cells. Each cell was annotated as malignant or non-malignant using scROSHI’s cell type annotations. Starting at the root node, we then iteratively assigned a CNV status to the nodes according to the composition of their subtrees. Specifically, a node and all nodes in its subtree were annotated as presenting no CNVs if both its subtrees contained at least 60% of non-malignant cells. We traversed the dendrogram until we reached all nodes or a maximum depth of five in the dendrogram. Finally, a cell was assigned the “no CNVs” status if it belonged to a leaf node that had been annotated as not presenting CNVs. All remaining cells were annotated as showing CNVs.

## Results & Discussion

### Performance evaluation

We compared the performance of scROSHI on test datasets with the performance of supervised methods that had been trained with the test dataset (intra-dataset evaluation) and that had been trained with a different dataset (inter-dataset evaluation). There were three types of classifiers: (1) prior knowledge method (scROSHI) for which a marker gene list is required. (2) Supervised methods (RF, SVM), which require a training dataset labeled with corresponding cell labels. (3) Combined method (GARNETT), which requires both a marker gene list and a training dataset. We calculated the percentage of unlabeled cells across all cell populations per classifier. Further, we calculated the accuracy of only major cell types for scROSHI and GARNETT, since both methods perform a hierarchical cell typing with major and subtype labels (Suppl. Tab. 5). Additionally, we determined the proportions of cells that only have a major cell type label, cells that have label “unknown”, or are unclassified.

Figure 2 shows the overall results of the inter and intra-dataset evaluation. Generally, scROSHI performs as well as the supervised methods if the supervised methods were trained with the test dataset (scROSHI accuracy: Adult 0.823, Newborn 0.879, Zheng 0.715). However, scROSHI outperforms the supervised methods if they were trained with another dataset - in this case we observed a lower accuracy and/or a higher amount of unlabeled cells for all supervised methods. The supervised methods perform better if they were trained with a dataset that is closer to the test dataset (e.g. training data: Adult; test data: Newborn) but there is a strong decrease in performance if the test data is dissimilar (e.g. training data: Adult; test data: Zheng).

**Figure 2:**
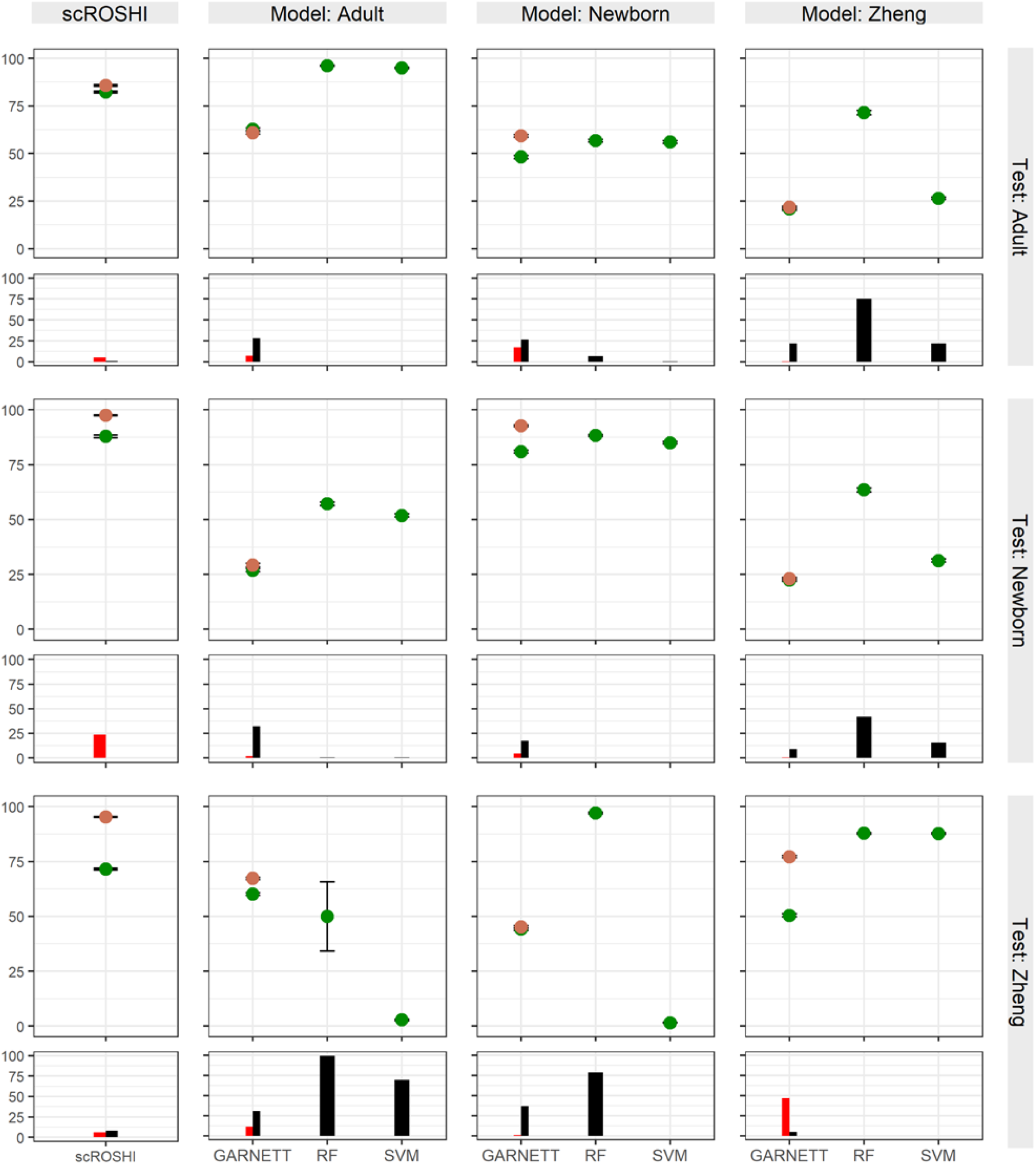
Benchmark results for scROSHI (left) and the three competing machine learning methods. Each panel corresponds to a combination of training data (row) and test data (column). The cross-validation accuracy is shown separately for major cell types (orange) and all fine-grained cell types (green). The red bar shows the proportion of cases where cells have only a major cell type label and subtype “unknown”. The black bar shows the proportion of cases where the cell type is “unknown”.

The subtype classification on the Zheng dataset was challenging for all classifiers (scROSHI accuracy: 0.715). However, the accuracy of the major cell type label was 0.952 for scROSHI indicating that even if it was not possible to find the correct subtype label the correct major cell type label could usually be determined.

We presented the design and validation of the robust cell type identification method scROSHI that performs superior to competing tools provided a good quality marker gene list is available and annotated training data are limited or not available, which is often a realistic scenario in early stage projects.

Similar to the scoring tool `ucell` ^28^, the cell type score of scROSHI depends only on the relative rank of the gene expression signal, does not require normalization, and makes no assumptions about the distribution of the signal. But, because scROSHI utilizes the hierarchical nature of cell identities, it can outperform its competitor when a sample contains similar cell types that derive from different branches of the lineage tree.

scROSHI was developed for 10xGenomics mRNAseq data of tumor patient samples but there is no known limitation to use it on any other modality or organism. However, it is ideal if the cell type-specific gene sets were defined from results of the same technology as the data at hand.

One possibility to improve the performance of the machine learning tools, i.e., the accuracy on unseen data, might be to train them on a more diverse data set. Yet, because training on accuracy does not learn causal features for cell type identity, this approach by design does not lead to a universally applicable model and the performance will still be low on unseen data.

### Consistency with estimations of copy number alterations

In addition to these benchmark datasets with known ground truth but relatively simple cell type composition we used scROSHI for cell type identification in clinical samples, i.e., biopsies from melanoma patients participating in the Tumor Profiler Study^20^. Without ground truth we evaluated the classification results by consistency with copy number variation (CNV) estimations (Fig. 2), and by comparison to single cell CyTOF cell type composition analysis on the same samples.

The three representative samples in Fig. 3A-C show a diverse cell type composition, as illustrated by the two-dimensional UMAP representation based on gene expression in the top row. CNV states appear nearly exclusively in cells identified as melanoma cells, the only malignant cells present (insets). In Fig. 3 bottom row, the focus is shifted to UMAP representations based on CNV states, where all non-malignant cells form a single cluster and malignant cells one or more separate clusters. In the sample shown in Fig. 3A, a few cells located in the melanoma cluster are mis-classified as cancer associated fibroblasts (CAFs, filled purple circles), possibly a consequence of an increased copy number in melanoma cells located at some CAF marker genes and/or copy number decrease in some melanoma marker genes.

**Figure 3:**
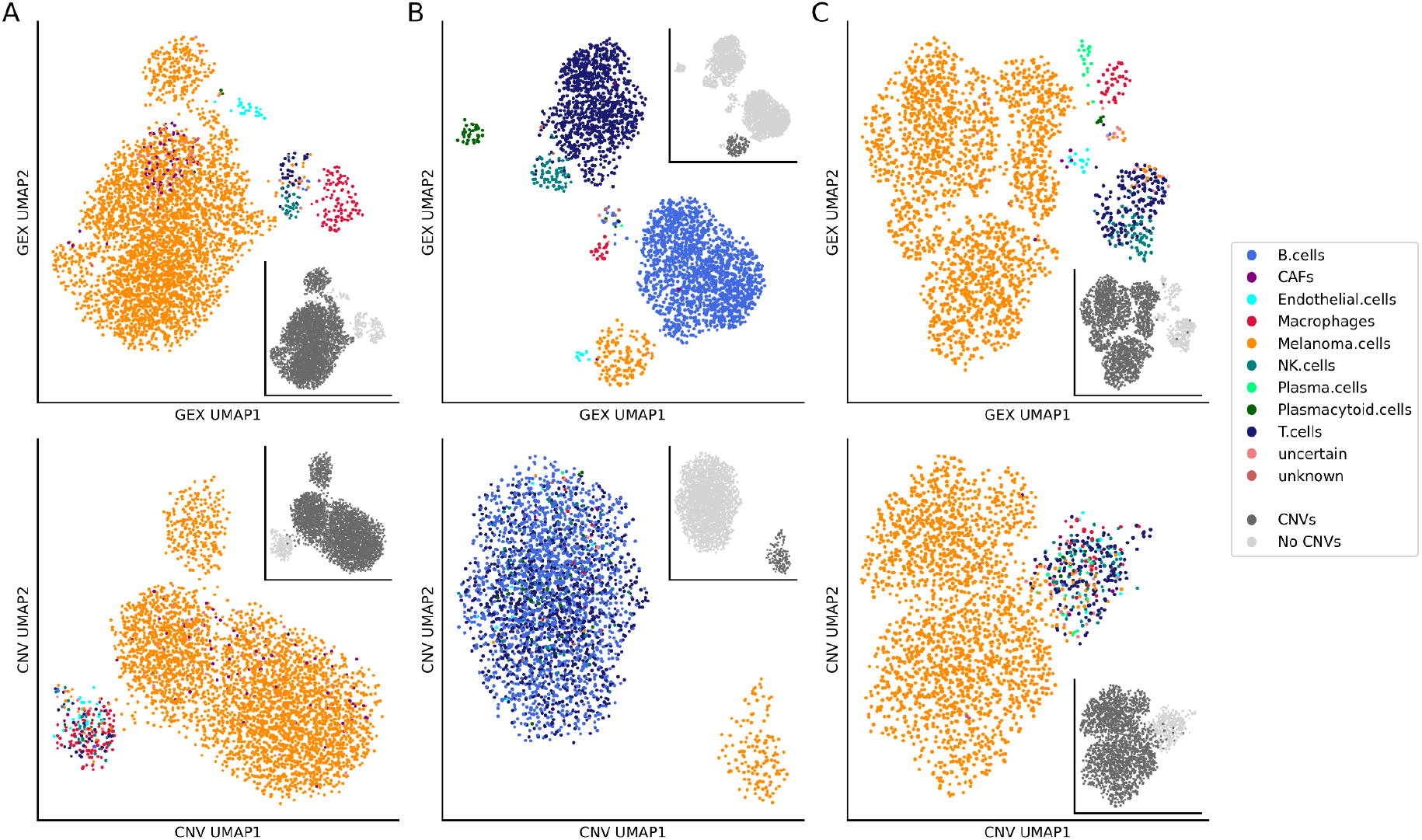
UMAP representation of cells in gene expression and CNV space. Cells from three melanoma biopsy samples (A n=3,928 cells; B n=2,967 cells; C n=2,326 cells) were annotated using scROSHI. The first row shows the UMAP embedding of normalized and log-transformed gene expression, the second row shows the UMAP embedding of CNV profiles. The colors represent the cell type annotations. The greyscale in the insets represent the CNV status.

The cell type composition in these samples is dominated by melanoma cells (A: 92%, B: 8%, C: 87%), B cells (A: 0%, B: 52%, C: 1%) and T cells (A: 1%, B: 33%, C: 7%). A comparison to single cell CyTOF experiments of the same samples gave similar proportions: melanoma cells (A: 81%, B: 2%, C: 84%), B cells (A: 0.7%, B: 40%, C: 2%) and T cells (A: 5%, B: 40%,C: 9%).

## Conclusion

Cell type identification is a critical, yet challenging, step in single cell transcriptomics analysis. Here, we have presented scROSHI, a novel supervised cell type classification method independent of training data but instead based on *a priori* defined cell type marker genes. We hope to have shown convincing evidence that scROSHI is useful, robust, versatile, and competitive to existing methods under real-life scenarios.

## Data Availability

Availability of benchmark data: ask at. The three data sets from the TumorProfiler Study are available upon request at, according to the data sharing policy at the web site https://eth-nexus.github.io/tu-pro_website/data/ (in preparation by the consortium).

Code availability: scROSHI will be available as R package on CRAN (in preparation). The source code for scROSHI is available on github at https://github.com/ETH-NEXUS/scROSHI.

## Acknowledgements

The authors would like to thank Ulrike Menzel (D-BSSE, ETHZ) for continuous support and Marcus Lindberg (NEXUS, ETHZ) for support in the early stages of the project. Further thanks are given to The Broad Institute of MIT and Harvard for early data access and use for benchmarking purposes.

