## Supplementary information for "scROSHI - robust supervised hierarchical identification of single cells"

### Supplementary Material

#### Section 1: Ground truth data sets

##### Data set Zheng_sorted

FASTQ files for the Zheng_sorted PBMC dataset [1] were retrieved from the '10X Genomics Single Cell Gene Expression Datasets' page at https://support.10xgenomics.com/single-cell-gene-expression/datasets - “Single Cell 3' Paper: Zheng et al. 2017”

The Zheng_sorted data set contains sequencing data derived from ten different cell types (refer to Suppl. Table 1), provided as separate fastq files. The different cell types have been individually mapped to the GRCh38 reference using *Cellranger count*, version 3.1 (reference version: refdata-cellranger-GRCh38-3.0.0; parameter setting --nosecondary). Afterwards, the output has been aggregated into a single data set using *Cellranger aggr.*

The original data set included 110526 cells, which we downsampled to 10% (11052) cells by taking 10% of the cells of each cell type.
The raw count data was further processed to remove low quality cells and to assign the respective ground truth label to each cell.

*Suppl. Table 1 : Zheng_sorted data set, cell numbers and cell type fractions*

| Zheng_sorted | | | | |
|---|---|---|---|---|
| Cell type | Raw - cell number | Raw - cell type fraction | Subset - cell number | Subset - cell type fraction |
| b_cells | 11257 | 10.2 | 1125 | 10.2 |
| cd14_monocytes | 11451 | 10.4 | 1145 | 10.4 |
| cd34 | 11680 | 10.6 | 1168 | 10.6 |
| cd4_t_helper | 11600 | 10.5 | 1160 | 10.5 |
| cd56_nk | 9110 | 8.2 | 911 | 8.2 |
| cytotoxic_t | 10511 | 9.5 | 1051 | 9.5 |
| memory_t | 10796 | 9.8 | 1080 | 9.8 |
| naive_cytotoxic | 12170 | 11 | 1217 | 11 |
| naive_t | 10758 | 9.7 | 1076 | 9.7 |
| regulatory_t | 11193 | 10.1 | 1119 | 10.1 |
| **Total** | **110526** | **100** | **11052** | **100** |

#### Data set Adult

2020-Mar-Census-Adult-Immune-10x dataset
The Adult PBMC data set was retrieved from the Broad Institute following a now disfunctional link https://singlecell.broadinstitute.org/single_cell/study/SCP771/2020-mar-census-adult-immune-10x. The same data is now annotated as "bone marrow" and available upon request at https://data.humancellatlas.org/explore/projects/cc95ff89-2e68-4a08-a234-480eca21ce79.
We would like to repeat our thanks to Aviv Regev and coworkers for their permission to use a small section of the data in the benchmark study and to publish it.

The Adult data set contained the count matrices already derived from cellranger count.
The original data set included 262579 cells from 16 different cell types, which we downsampled to 10% (26250) cells by taking 10% of the cells of each cell type.

*Suppl Table 2 : Adult data set, cell numbers and cell type fractions*

| Adult | | | | |
|---|---|---|---|---|
| Cell type | Raw - cell number | Raw - cell type fraction | Subset - cell number | Subset - cell type fraction |
| CD14+ monocyte type 1 | 37086 | 14.1 | 3708 | 14.1 |
| CD14+ monocyte type 2 | 7151 | 2.7 | 715 | 2.7 |
| CD16+ monocyte | 4065 | 1.5 | 406 | 1.5 |
| CD4+ naive T cell | 49169 | 18.7 | 4916 | 18.7 |
| conventional dendritic cell | 4494 | 1.7 | 449 | 1.7 |
| cytotoxic T cell type 1 | 22336 | 8.5 | 2233 | 8.5 |
| cytotoxic T cell type 2 | 15740 | 6 | 1574 | 6 |
| memory B cell | 9139 | 3.5 | 913 | 3.5 |
| naive B cell | 26547 | 10.1 | 2654 | 10.1 |
| naive CD8+ T cell | 20464 | 7.8 | 2046 | 7.8 |
| natural killer cell | 20554 | 7.8 | 2055 | 7.8 |
| plasma cell | 2687 | 1 | 268 | 1 |
| plasmacytoid dendritic cell | 2735 | 1 | 273 | 1 |
| precursor B cell | 8200 | 3.1 | 820 | 3.1 |
| pro-B cell | 3398 | 1.3 | 339 | 1.3 |
| T-helper cell | 28814 | 11 | 2881 | 11 |
| **Total** | **262579** | **100** | **26250** | **100** |

#### Data set Newborn

2020-Mar-Census-Newborn-Immune-10x dataset
The Newborn PBMC data set was retrieved from the Broad Institute following a now disfunctional link:
https://singlecell.broadinstitute.org/single_cell/study/SCP770/2020-mar-census-newborn-blood-10x
The same data is now annotated as “umbilical cord blood” and available upon request at https://data.humancellatlas.org/explore/projects/cc95ff89-2e68-4a08-a234-480eca21ce79.
We would like to repeat our thanks to Aviv Regev and coworkers for their permission to use a small section of the data in the benchmark study and to publish it.

The Newborn data set contained the count matrices already derived from cellranger count. The original data set included 196278 cells from 12 different cell types, which we downsampled to 10% of cells by taking 10% of the cells of each cell type. Further, we excluded the cell type “T-helper cell including regulatory T cell” as this did not specify a clearly defined single cell type but a mix of types. Overall, the Newborn set was reduced to 18887 cells.

*Suppl Table 3 : Newborn data set, cell numbers and cell type fractions*

| Newborn | | | | |
|---|---|---|---|---|
| Cell type | Raw - cell number | Raw - cell type fraction | Subset - cell number | Subset - cell type fraction |
| CD14+ monocyte type 1 | 27274 | 13.9 | 2727 | 14.4 |
| CD14+ monocyte type 2 | 1580 | 0.8 | 158 | 0.8 |
| CD4+ T cell | 2514 | 1.3 | 251 | 1.3 |
| dendritic cell | 1665 | 0.8 | 166 | 0.9 |
| memory B cell | 1692 | 0.9 | 169 | 0.9 |
| naive B cell | 36441 | 18.6 | 3644 | 19.3 |
| naive CD8+ T cell | 33593 | 17.1 | 3359 | 17.8 |
| naive T-helper cell type 1 | 36795 | 18.7 | 3679 | 19.5 |
| naive T-helper cell type 2 | 32690 | 16.7 | 3269 | 17.3 |
| natural killer cell type 1 | 12696 | 6.5 | 1269 | 6.7 |
| natural killer cell type 2 | 1962 | 1 | 196 | 1 |
| T-helper cell including regulatory T cell | 7374 | 3.8 | 0 | 0 |
| **Total** | **196276** | **100** | **18887** | **100** |

### Section 2: Ground truth dictionaries

#### Cell type label harmonization

To accommodate comparisons between data sets, we defined a common set of cell type labels matching between the three different ground truth data sets (refer to Suppl Table 4). In order to harmonize cell type labels based on visual inspection we combined the three data sets and visualized the cell types together in a UMAP.
As is shown in Suppl Figures 1-3, the embedding is dominated by the data set differences rather than cell type differences. Nevertheless, groups of cell type populations can be observed, mainly a B cell cluster, a T cell cluster, and a monocyte and stem cell cluster.
When cells of differently labeled cell types were in close proximity in the UMAP embedding, we chose the same label for both types.

Regarding B cell sub types (visualized in Suppl Fig 1), only very few Memory B cells can be observed for the Newborn data set, and moreover they are grouped together with the Naive B cells. The same can be observed for the Adult set, also here Memory B cells and Naive B cells group together. Only the Precursor B cells cluster separately from the other B cell sub types. Accordingly, we merged the sub type Memory B cell into Naive B cell.

Regarding the Monocyte and Stem cell sub types (visualized in Suppl Fig 2), we again observe the data set identity as the major driver for the distinct cell type populations. Of note, for the Zheng_sorted data set the Monocytes group together with the stem cells, and further the stem cells show diverse sub populations. This already indicates that it will be difficult to use Zheng_sorted Monocytes to type Adult and Newborn Monocytes. The Monocyte sub types from the Adult set show very distinct clusters and are also very close to the Dendritic cells. The same holds true for the Newborn set. Here, the CD14 Monocyte Type 2 is also a very small set that overlaps with the CD14 Monocyte Type 1 cells. Taken together, we harmonized cell type labels by merging the different Monocyte sub types into one major class called Monocytes.
The few B cells and T cells presented in this UMAP are potentially doublets or misclassified cells, because they do not cluster with the rest of their respective cell type.

Regarding the T cell and NK cell sub types (visualized in Supp Fig 3), it is notable that for the Zheng_sorted data set all T cell sub types are spread out without forming a distinct population per sub type. The only exception are the NK cells, where the majority of cells forms one cluster. Further, for the Adult data set Naive CD4 positive and Naive CD8 positive T cells are completely overlapping each other. This indicates that for cell typing it will be very difficult to distinguish both types. The same holds true for the Naive T-helper cell Type 1 and Type 2 and CD4 positive T cell sub types. They are very similar and thus it is doubtful whether an expression-based cell type classification could distinguish these sub types.
Taken together, we harmonized the cell type labels and e.g. grouped Naive T helper cells into CD4 T cells, Cytotoxic CD8 sub types into CD8 T cells.
The details are shown in Suppl Table 4.

*Suppl Figure 1: UMAP representation of the B cell sub types present across the three PBMC ground truth data sets. Memory B cells and Naive B cells are very similar to each other, for Newborn and Adult, respectively. The population distances are driven by data set, not by cell type.*

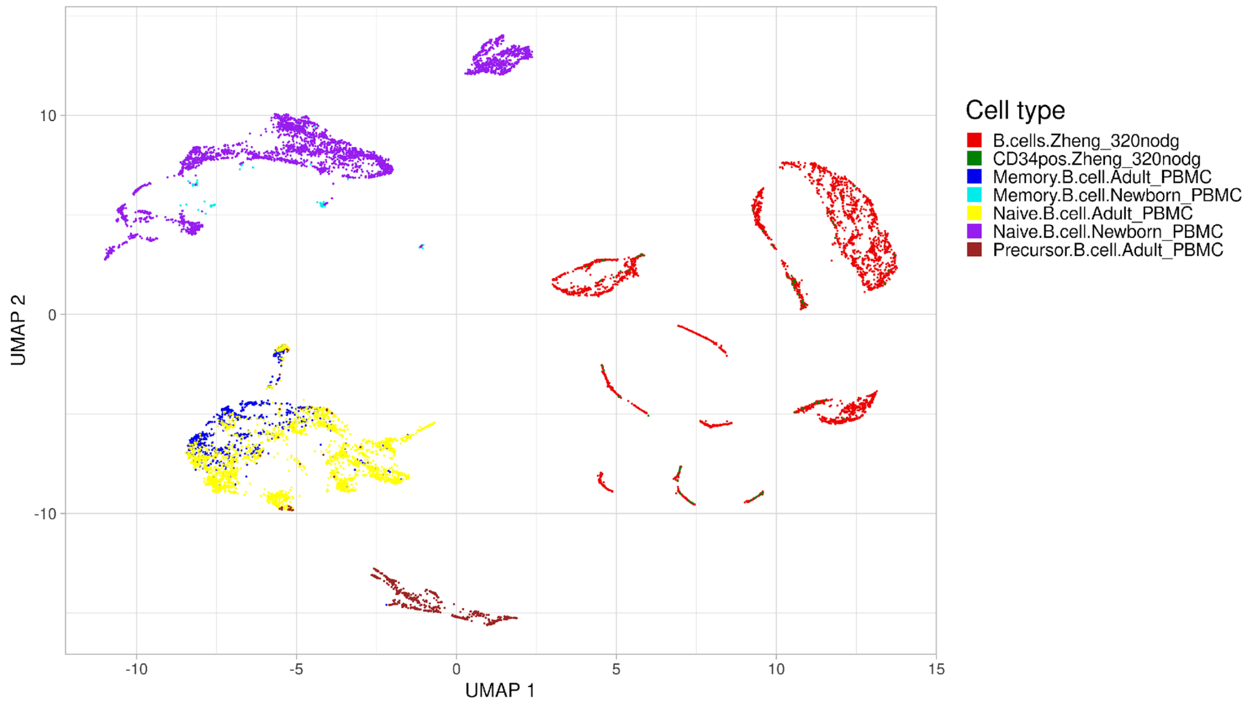


*Suppl Figure 2: UMAP representation of the Monocyte and stem cell types present across the three PBMC ground truth data sets. Note that very few B cells and T cells are present as well, as they grouped closer with the Monocyte sub types than with the rest of the B cells and T cells.*

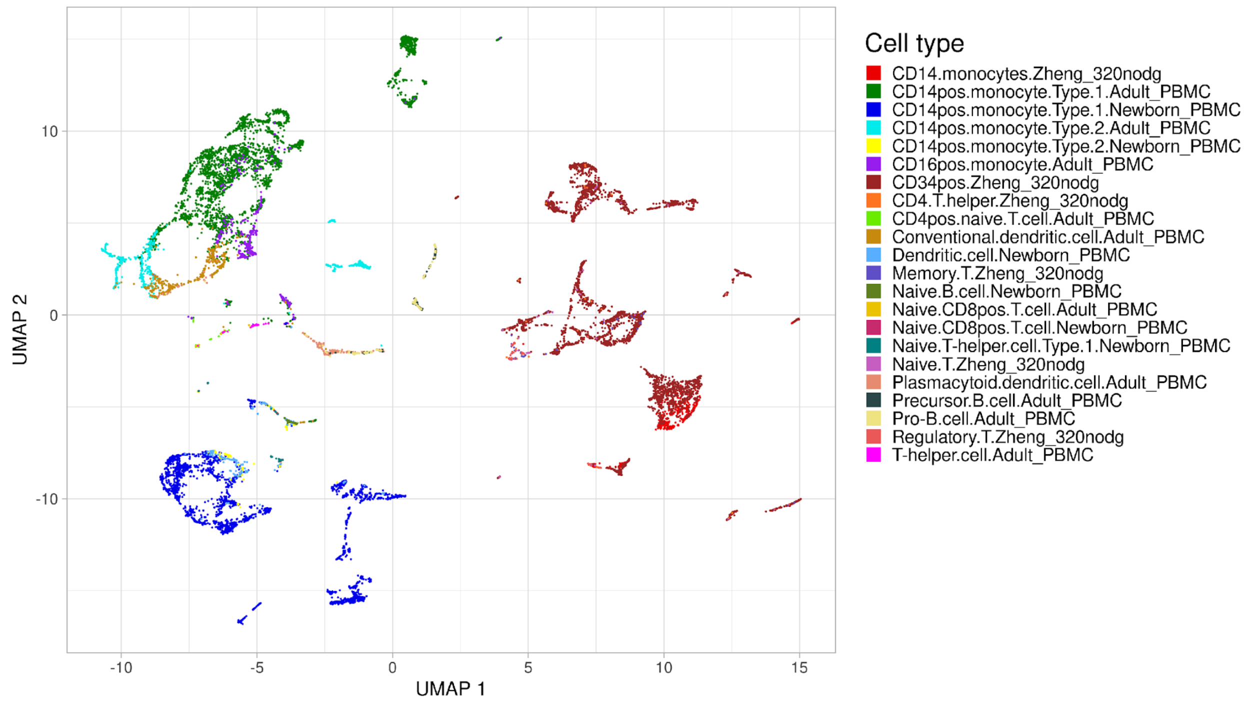

*Suppl Figure 3: UMAP representation of the T cell and NK cell sub types present across the three PBMC ground truth data sets.*

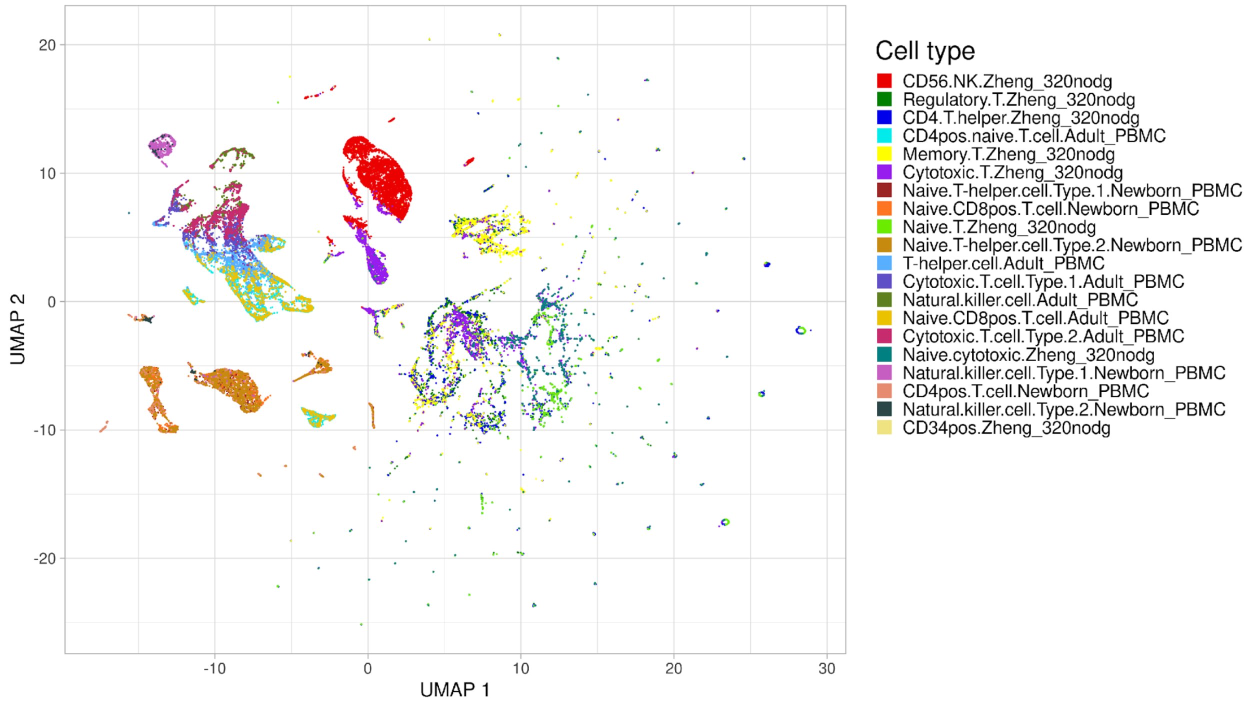

*Suppl Table 4: Ground truth labels for PBMC data sets*

| Original | Matched | Data set |
|---|---|---|
| b_cells | B.cells | Zheng_sorted |
| cd14_monocytes | Monocytes | Zheng_sorted |
| cd34 | Stem.cells | Zheng_sorted |
| cd4_t_helper | CD4.T.cells | Zheng_sorted |
| cd56_nk | NK.cells | Zheng_sorted |
| cytotoxic_t | CD8.T.cells | Zheng_sorted |
| naive_cytotoxic | CD8.T.cells | Zheng_sorted |
| regulatory_t | T.reg.cells | Zheng_sorted |
| memory_t | CD4.T.cells | Zheng_sorted |
| naive_t | CD4.T.cells | Zheng_sorted |
| CD14+ monocyte type 1 | Monocytes | Adult |
| CD14+ monocyte type 2 | Monocytes | Adult |
| CD16+ monocyte | Monocytes | Adult |
| CD4+ naive T cell | CD4.T.cells | Adult |
| conventional dendritic cell | Dendritic.cells | Adult |
| cytotoxic T cell type 1 | CD8.T.cells | Adult |
| cytotoxic T cell type 2 | CD8.T.cells | Adult |
| memory B cell | B.cells.naive | Adult |
| naive B cell | B.cells.naive | Adult |
| naive CD8+ T cell | CD4.T.cells | Adult |
| natural killer cell | NK.cells | Adult |
| plasma cell | Plasma.cells | Adult |
| plasmacytoid dendritic cell | Plasmacytoid.dendritic.cells | Adult |
| precursor B cell | B.cells.precursor | Adult |
| pro-B cell | B.cells.precursor | Adult |
| T-helper cell | CD4.T.cells | Adult |
| CD14+ monocyte type 1 | Monocytes | Newborn |
| CD14+ monocyte type 2 | Monocytes | Newborn |
| CD4+ T cell | CD4.T.cells | Newborn |
| dendritic cell | Dendritic.cells | Newborn |
| memory B cell | B.cells.naive | Newborn |
| naive B cell | B.cells.naive | Newborn |
| naive CD8+ T cell | CD8.T.cells | Newborn |
| naive T-helper cell type 1 | CD4.T.cells | Newborn |
| naive T-helper cell type 2 | CD4.T.cells | Newborn |
| natural killer cell type 1 | NK.cells | Newborn |
| natural killer cell type 2 | NK.cells | Newborn |

##### Hierarchical cell typing:

scROSHI and Garnett are able to perform a hierarchical cell type classification to first distinguish into major lineage classes (e.g. T cell versus Endothelial cell) and further into sub types (e.g. into different T cell sub types).
To accommodate this hierarchical cell typing, we grouped the ground truth cell types according to their lineage into *Major* and *Subtype* classes (see suppl Table 5). These groups specified the config files necessary to run Garnett and scROSHI.

*Suppl. Table 5: Hierarchical cell type classes*

| **Major_type** | **Subtypes** |
|---|---|
| Monocytes | none |
| T.cells | T.cells.CD4, T.cells.CD8, T.cells.regulatory |
| Dendritic.cells | none |
| B.cells | B.cells.naive, B.cells.precursor |
| Plasma.cells | none |
| Plasmacytoid.dendritic.cells | none |
| NK.cells | none |
| Stem.cells | none |

#### Section 3: Benchmark pipeline

We implemented a snakemake-based [2] benchmark pipeline to accommodate the processing of the three different ground truth PBMC data sets and to allow a seamless integration of new benchmark data sets in the future.
The framework is available on github (https://github.com/ETH-NEXUS/cell_type_benchmark.git) and comprises two main workflows: *training* and *testing*.
The *training* workflow takes as input bioconductor single cell experiment objects (SCE) of the different data sets[3]. The SCE needs to contain the gene x cell count matrix as well as the ground truth label of each cell. The repository contains the pre-processing scripts applied to subset the benchmark data sets and to add ground-truth cell type labels to the SCE objects.
The *training* workflow facilitates (i) the cell type marker classification based on Seurat for each of the input data sets (necessary e.g. for tools like Garnett and scROSHI that need input marker files). Second, it performs a cross validation and model training for the cell typing methods that need a pre-trained model for classification (e.g. SVM, RF, and Garnett).
The trained model as well as the cell type marker file is then passed on to the *testing* workflow. Here, for each data set combination (e.g. trained on Adult and tested on Newborn) the cell typing methods are applied and the result is formatted to allow the evaluation framework to be applied.

#### Evaluation framework

The overall accuracy is reported as a measurement for the performance on the dataset. Additionally, a 95 percent confidence interval was computed using the R function *binom.test()*. The used methods do not label all cells (unknown). These unknown cells are not considered in the accuracy calculations. The percentage of unlabeled cells is also used to evaluate the performance. We determined the accuracy for the major cell type separately for the hierarchical classification methods (scROSHI and GARNETT) along with the fraction of cells that only have a major cell type label and no subtype label. Furthermore, we calculated F1 statistics for each cell type and each method. The F1 statistics for the major cell types were determined in addition to the hierarchical methods.

### TumorProfiler Consortium

Rudolf Aebersold[2], Melike Ak[27], Faisal S Al-Quaddoomi[9,16], Jonas Albinus[7], Ilaria Alborelli[23], Sonali Andani[6,16,25,30], Per-Olof Attinger[11], Marina Bacac[15], Daniel Baumhoer[23], Beatrice Beck-Schimmer[38], Niko Beerenwinkel[4,16], Christian Beisel[4], Lara Bernasconi[26], Anne Bertolini[9,16], Bernd Bodenmiller[8,34], Ximena Bonilla[6,16,25], Lars Bosshard[9,16], Byron Calgua[23], Ruben Casanova[34], Stéphane Chevrier[34], Natalia Chicherova[9,16], Maya D'Costa[10], Esther Danenberg[36], Natalie Davidson[6,16,25], Monica-Andreea Drăgan[4], Reinhard Dummer[27], Stefanie Engler[34], Martin Erkens[13], Katja Eschbach[4], Cinzia Esposito[36], André Fedier[17], Pedro Ferreira[4], Joanna Ficek[6,16,25], Anja L Frei[30], Bruno Frey[12], Sandra Goetze[7], Linda Grob[9,16], Gabriele Gut[36], Detlef Günther[5], Martina Haberecker[30], Pirmin Haeuptle[1], Viola Heinzelmann-Schwarz[17,22], Sylvia Herter[15], Rene Holtackers[36], Tamara Huesser[15], Anja Irmisch[27], Francis Jacob[17], Andrea Jacobs[34], Tim M Jaeger[11], Katharina Jahn[4], Alva R James[6,16,25], Philip M Jermann[23], André Kahles[6,16,25], Abdullah Kahraman[16,30], Viktor H Koelzer[30], Werner Kuebler[24], Jack Kuipers[4,16], Christian P Kunze[21], Christian Kurzeder[20], Kjong-Van Lehmann[6,16,25], Mitchell Levesque[27], Sebastian Lugert[10], Gerd Maass[12], Markus G Manz[29], Philipp Markolin[6,16,25], Julien Mena[2], Ulrike Menzel[4], Julian M Metzler[28], Nicola Miglino[1], Emanuela S Milani[7], Holger Moch[30], Simone Muenst[23], Riccardo Murri[37], Charlotte KY Ng[23,33], Stefan Nicolet[23], Marta Nowak[30], Patrick GA Pedrioli[3], Lucas Pelkmans[36], Salvatore Piscuoglio[17,23], Michael Prummer[9,16], Mathilde Ritter[17], Christian Rommel[13], María L Rosano-González[9,16], Gunnar Rätsch[3,6,16,25], Natascha Santacroce[4], Jacobo Sarabia del Castillo[36], Ramona Schlenker[14], Petra C Schwalie[13], Severin Schwan[11], Tobias Schär[4], Gabriela Senti[26], Franziska Singer[9,16], Sujana Sivapatham[34], Berend Snijder[2,16], Bettina Sobottka[30], Vipin T Sreedharan[9,16], Stefan Stark[6,16,25], Daniel J Stekhoven[9,16], Alexandre PA Theocharides[29], Tinu M Thomas[6,16,25], Markus Tolnay[23], Vinko Tosevski[15], Nora C Toussaint[9,16], Mustafa A Tuncel[4,16], Marina Tusup[27], Audrey Van Drogen[7], Marcus Vetter[19], Tatjana Vlajnic[23], Sandra Weber[26], Walter P Weber[18], Rebekka Wegmann[2], Michael Weller[32], Fabian Wendt[7], Norbert Wey[30], Andreas Wicki[29,35], Mattheus HE Wildschut[2,29], Bernd Wollscheid[7], Shuqing Yu[9,16], Johanna Ziegler[27], Marc Zimmermann[6,16,25], Martin Zoche[30], Gregor Zuend[31]

[1]Cantonal Hospital Baselland, Medical University Clinic, Rheinstrasse 26, 4410 Liestal, Switzerland, [2]ETH Zurich, Department of Biology, Institute of Molecular Systems Biology, Otto-Stern-Weg 3, 8093 Zurich, Switzerland, [3]ETH Zurich, Department of Biology, Wolfgang-Pauli-Strasse 27, 8093 Zurich, Switzerland, [4]ETH Zurich, Department of Biosystems Science and Engineering, Mattenstrasse 26, 4058 Basel, Switzerland, [5]ETH Zurich, Department of Chemistry and Applied Biosciences, Vladimir-Prelog-Weg 1-5/10, 8093 Zurich, Switzerland, [6]ETH Zurich, Department of Computer Science, Institute of Machine Learning, Universitätstrasse 6, 8092 Zurich, Switzerland, [7]ETH Zurich, Department of Health Sciences and Technology, Otto-Stern-Weg 3, 8093 Zurich, Switzerland, [8]ETH Zurich, Institute of Molecular Health Sciences, Otto-Stern-Weg 7, 8093 Zurich, Switzerland, [9]ETH Zurich, NEXUS Personalized Health Technologies, John-von-Neumann-Weg 9, 8093 Zurich, Switzerland, [10]F. Hoffmann-La Roche Ltd, Grenzacherstrasse 124, 4070 Basel, Switzerland, [11]F. Hoffmann-La Roche Ltd, Grenzacherstrasse 124, 4070 Basel, Switzerland, , [12]Roche Diagnostics GmbH, Nonnenwald 2, 82377 Penzberg, Germany, [13]Roche Pharmaceutical Research and Early Development, Roche Innovation Center Basel, Grenzacherstrasse 124, 4070 Basel, Switzerland, [14]Roche Pharmaceutical Research and

Early Development, Roche Innovation Center Munich, Roche Diagnostics GmbH, Nonnenwald 2, 82377 Penzberg, Germany, [15]Roche Pharmaceutical Research and Early Development, Roche Innovation Center Zurich, Wagistrasse 10, 8952 Schlieren, Switzerland, [16]SIB Swiss Institute of Bioinformatics, Lausanne, Switzerland, [17]University Hospital Basel and University of Basel, Department of Biomedicine, Hebelstrasse 20, 4031 Basel, Switzerland, [18]University Hospital Basel and University of Basel, Department of Surgery, Brustzentrum, Spitalstrasse 21, 4031 Basel, Switzerland, [19]University Hospital Basel, Brustzentrum & Tumorzentrum, Petersgraben 4, 4031 Basel, Switzerland, [20]University Hospital Basel, Brustzentrum, Spitalstrasse 21, 4031 Basel, Switzerland, [21]University Hospital Basel, Department of Information- and Communication Technology, Spitalstrasse 26, 4031 Basel, Switzerland, [22]University Hospital Basel, Gynecological Cancer Center, Spitalstrasse 21, 4031 Basel, Switzerland, [23]University Hospital Basel, Institute of Medical Genetics and Pathology, Schönbeinstrasse 40, 4031 Basel, Switzerland, [24]University Hospital Basel, Spitalstrasse 21/Petersgraben 4, 4031 Basel, Switzerland, [25]University Hospital Zurich, Biomedical Informatics, Schmelzbergstrasse 26, 8006 Zurich, Switzerland, [26]University Hospital Zurich, Clinical Trials Center, Rämistrasse 100, 8091 Zurich, Switzerland, [27]University Hospital Zurich, Department of Dermatology, Gloriastrasse 31, 8091 Zurich, Switzerland, [28]University Hospital Zurich, Department of Gynecology, Frauenklinikstrasse 10, 8091 Zurich, Switzerland, [29]University Hospital Zurich, Department of Medical Oncology and Hematology, Rämistrasse 100, 8091 Zurich, Switzerland, [30]University Hospital Zurich, Department of Pathology and Molecular Pathology, Schmelzbergstrasse 12, 8091 Zurich, Switzerland, [31]University Hospital Zurich, Rämistrasse 100, 8091 Zurich, Switzerland, [32]University Hospital and University of Zurich, Department of Neurology, Frauenklinikstrasse 26, 8091 Zurich, Switzerland, [33]University of Bern, Department of BioMedical Research, Murtenstrasse 35, 3008 Bern, Switzerland, [34]University of Zurich, Department of Quantitative Biomedicine, Winterthurerstrasse 190, 8057 Zurich, Switzerland, [35]University of Zurich, Faculty of Medicine, Zurich, Switzerland, [36]University of Zurich, Institute of Molecular Life Sciences, Winterthurerstrasse 190, 8057 Zurich, Switzerland, [37]University of Zurich, Services and Support for Science IT, Winterthurerstrasse 190, 8057 Zurich, Switzerland, [38]University of Zurich, VP Medicine, Künstlergasse 15, 8001 Zurich, Switzerland
